# Ligand Modulates Cross-Coupling between Riboswitch Folding and Transcriptional Pausing

**DOI:** 10.1101/350447

**Authors:** Julia R. Widom, Yuri A. Nedialkov, Victoria Rai, Ryan L. Hayes, Charles L. Brooks, Irina Artsimovitch, Nils G. Walter

## Abstract

**Highlights:**

- smFRET, biochemistry and simulations probe co-transcriptional riboswitch folding
- Nascent RNA folding is disfavored by the DNA template and aided by RNAP
- Transcriptional pausing is stabilized by RNA pseudoknot and destabilized by ligand

**SUMMARY:** Numerous classes of riboswitches have been found to regulate bacterial gene expression in response to physiological cues, offering new paths to anti-bacterial drugs. As common studies of isolated riboswitches lack the functional context of the transcription machinery, we here combine single-molecule, biochemical and simulation approaches to investigate the coupling between co-transcriptional folding of the pseudoknot-structured preQ_1_ riboswitch and RNA polymerase (RNAP) pausing. We show that pausing at a site immediately downstream of the riboswitch requires a ligand-free pseudoknot in the nascent RNA, a precisely spaced sequence resembling the pause consensus, and electrostatic and steric interactions with the RNAP exit channel. While interactions with RNAP stabilize the native fold of the riboswitch, binding of the ligand signals RNAP release from the pause. Our results demonstrate that the nascent riboswitch and its ligand actively modulate the function of RNAP and vice versa, a paradigm likely to apply to other cellular RNA transcripts.

## INTRODUCTION

To render bacteria responsive to rapidly changing environmental conditions, gene expression is regulated by a variety of means, including mechanisms in which riboswitches modulate messenger RNA (mRNA) synthesis, translation or stability in response to physiological cues. Riboswitches, structured non-coding RNA elements typically found in 5’ untranslated regions, are thought to regulate the expression of up to 4% of genes in certain bacteria (Sherwood and Henkin, 2016). A typical riboswitch contains a ligand-binding aptamer domain upstream of an expression platform whose conformation changes in response to ligand binding by the aptamer domain, thereby regulating expression of the downstream gene. The at least 40 classes of riboswitches for which ligands have been identified thus far (Greenlee *et al.*, 2018) sense diverse ligands including metabolites (Roth *et al.*, 2007), nucleobases (Frieda and Block, 2012), cofactors (Johnson *et al.*, 2012), other RNAs (Zhang and Ferré-D’Amaré, 2013), and metal ions (Dambach *et al.*, 2015). Due to their critical importance in bacterial gene regulation, riboswitches present attractive targets for the development of new antibiotic therapies (Deigan and Ferré-D’Amaré, 2011).

Mechanistic studies have been performed on many isolated riboswitches (Duesterberg *et al.*, 2015; Holmstrom *et al.*, 2014; Liberman *et al.*, 2015; Suddala *et al.*, 2013); however, in the cell, riboswitch folding occurs in the context of the transcription machinery. Two critical features are imposed on riboswitch folding as a result of its co-transcriptional nature: hierarchical folding in the 5’-to-3’ direction, and kinetic regulation by the varying speed of transcription, which places time limits on ligand sensing if the riboswitch is to regulate the outcome of transcription (Frieda and Block, 2012; Lutz *et al.*, 2014; Wickiser *et al.*, 2005). Conversely, riboswitch folding itself has the potential to modify transcription speed, but few examples of direct cross-talk between nascent RNA folding and transcription have been investigated. Most studies of co-transcriptional RNA folding utilize readouts that result from transcription itself, including assessing the activity of transcribed ribozymes (Pan *et al.*, 1999), RNase H probing to detect the accessibility of specific nucleotides (Chauvier *et al.*, 2017), SHAPE probing to detect backbone flexibility (Watters *et al.*, 2016), and single-molecule force and fluorescence techniques (Frieda and Block, 2012; Uhm *et al.*, 2018). When naked RNA is compared to RNA folded during active transcription, the effects of interactions with the transcription machinery are convoluted with the kinetic effects of non-equilibrium folding. As a result, potential direct effects of the presence of the DNA template and RNAP on folding remain largely unexplored.

The preQ_1_ riboswitch from the Gram-positive bacterium *Bacillus subtilis* (*Bsu*) is thought to act through a transcriptional mechanism in which the binding of its ligand, the tRNA nucleotide precursor 7-methylamino-7-deazaguanine (preQ_1_), stabilizes a pseudoknot structure that favors the formation of a terminator hairpin, decreasing expression of enzymes involved in queuosine biosynthesis (Figure 1A) (Roth *et al.*, 2007). With a minimal aptamer containing only 36 nucleotides, it is one of the smallest known riboswitches. By investigating this riboswitch in its greater biological context, we show that a network of interactions between the nascent RNA, DNA template and RNAP enable the transcription machinery to cross-couple regulation of nascent RNA folding and transcriptional pausing. Specifically, we used single-molecule FRET (smFRET) to probe the conformation of the nascent riboswitch at and beyond a transcriptional pause observed immediately after the aptamer domain is synthesized (U46, Figure S1). We found that, compared to the isolated RNA, the addition of a DNA scaffold that mimics a transcription bubble disfavors folding of the riboswitch, while the subsequent addition of RNAP recovers folding. Additionally, we found that the U46 pause is stabilized by interactions between a nascent RNA pseudoknot and RNAP in conjunction with a precisely spaced sequence that resembles the pause consensus identified *in vivo* (Figure S1D) (Larson *et al.*, 2014; Vvedenskaya *et al.*, 2014), and is destabilized (released) upon ligand binding. This pause, hereafter called the “*que* pause”, exhibits features distinct from the two previously defined mechanisms for transcriptional pausing in bacteria (Artsimovitch and Landick, 2000), leading us to term it “Class III”. Our observations demonstrate an unprecedented cross-coupling between riboswitch folding and RNAP pausing that is likely to govern the co-transcriptional folding of numerous RNAs.

**Figure 1.**
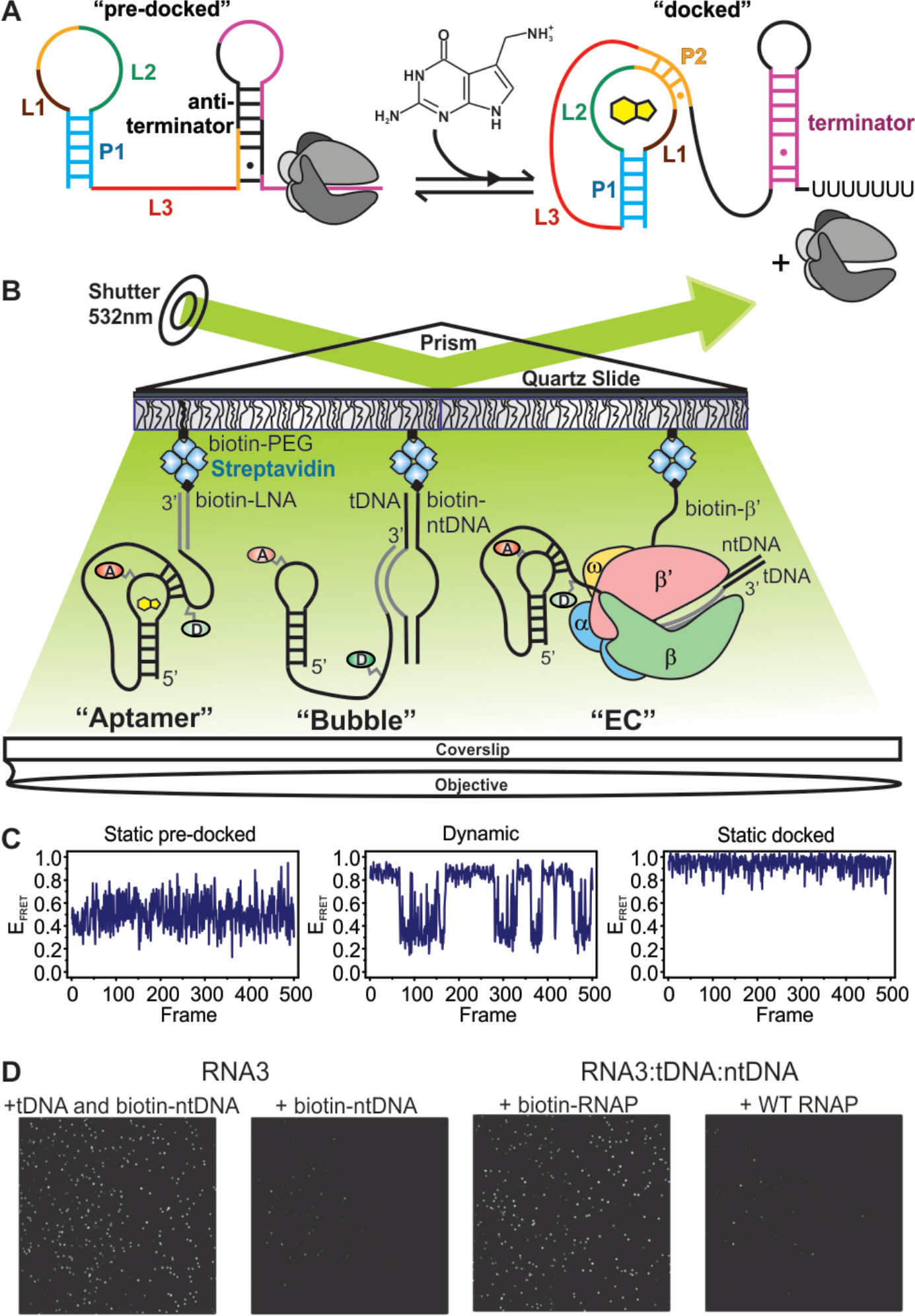
smFRET Investigation of the preQi Riboswitch in Active Transcription Complexes. (A) Regulation of gene expression by the preQ_1_ riboswitch. Ligand-induced pseudoknot docking favors the formation of a terminator hairpin, leading to decreased expression of the downstream genes. (B) smFRET experimental setup. “Aptamer” immobilization utilizes a biotinylated LNA capture probe. “Bubble” immobilization utilizes a partially complementary DNA bubble with a biotinylated nontemplate DNA. Elongation complex (“EC”) immobilization utilizes biotinylated *Eco* RNAP. Locations of donor (“D”, green) and acceptor (“A”, red) fluorophores are indicated. (C) Representative smFRET traces illustrating the three types of typical behavior observed across all samples. (D) iCCD images show that bubble immobilization is dependent on the tDNA in addition to the biotinylated ntDNA and the fluorophore-labeled RNA (left). Efficient EC immobilization requires biotinylated RNAP (right).

## RESULTS

### Direct Observation of Riboswitch Folding in Transcription Complexes

Due to the sensitivity of RNA folding to macromolecular crowding (Daher *et al.*, 2018; Dupuis *et al.*, 2014), ionic conditions (Suddala *et al.*, 2015) and other factors, we hypothesized that the behavior of the preQ_1_ riboswitch would be significantly altered upon incorporation into a transcription elongation complex (EC) containing a DNA template and RNAP. We therefore used smFRET to investigate the effects of the transcription machinery on the folding of the nascent riboswitch. We immobilized the riboswitch on polyethylene glycol (PEG)-passivated, streptavidin-coated quartz slides for imaging via prism-based total internal reflection fluorescence (p-TIRF) microscopy. The donor fluorophore was placed at the 2’ position of a guanosine residue within the L3 “tail” of the riboswitch, and the acceptor was placed on a 4-aminoallyl-uracil residue within the loop L2 (Figure 1). In an isolated RNA aptamer, the “docked” pseudoknot conformation in which the helix P2 is formed yields a high FRET efficiency (*E*_*FRET*_), while a mid-FRET state is observed for the “pre-docked” conformation in which P2 is not fully intact (Santner *et al.*, 2012; Suddala *et al.*, 2013; 2015) (Figure 1C).

We used a systematic series of immobilization strategies to study the riboswitch at varying degrees of biological complexity (Figure 1B). Immobilization through a biotinylated LNA capture probe (CP) allowed investigation of the isolated aptamer (henceforth called “aptamer immobilization”). A partially complementary DNA bubble with a biotinylated non-template (nt) strand enabled investigation of the nucleic acid framework of an EC (“bubble immobilization”). Finally, immobilization through biotinylated *Escherichia coli* (*Eco*) RNAP allowed observation of the riboswitch in a complete EC (“EC immobilization”) (Daube and Hippel, 1992; Sidorenkov *et al.*, 1998). Transcription by *Eco* and *Bsu* RNAPs yielded comparable results, justifying the use of the better studied *Eco* RNAP in this study, which sought to identify not only specific features of preQ_1_ riboswitch folding in the context of a known transcription machinery, but also factors that may contribute to the folding of nascent RNA in general (Figure S1).

To investigate co-transcriptional folding wherein riboswitch properties and interactions with the transcription machinery may evolve as additional RNA is synthesized, we utilized RNAs containing varying amounts of the riboswitch expression platform. The shortest RNA species studied by smFRET is analogous to the species present at the *que* pause (“RNA0 pause”, henceforth called “RNA0p”); we also investigated RNAs with an additional three (RNA3) or ten (RNA10) nucleotides of expression platform (Figure S2A) We first confirmed that our assembly procedures yielded the desired complexes and immobilization mechanisms. Efficient bubble immobilization required that the template DNA (tDNA) be present along with the fluorophore-labeled RNA and biotinylated ntDNA, indicating that the molecules we observe contain all three components. EC immobilization was efficient only with biotinylated RNAP, indicating the absence of nonspecific sticking to the slide (Figure 1D).

### Addition of the DNA Template Electrostatically Disfavors Riboswitch Folding

Aptamer-immobilized complexes were expected to exhibit behavior consistent with previous studies of the isolated riboswitch aptamer (Suddala *et al.*, 2013; 2015). Accordingly, we observed that in the absence of Mg^2+^ or preQ_1_, RNA0p existed almost exclusively in the pre-docked mid-FRET state with *E*_*FRET*_ ~ 0.5, while RNA3 also exhibited a minor fully-docked high-FRET population (15%, *E*_*FRET*_ = 0.88) (Figure 2A). For both RNAs, addition of Mg^2^+ shifted the mid-FRET state to *E*_*FRET*_ ~ 0.7, indicative of a more compact pre-docked conformation, while increasing the population of the high-FRET state (Figure 3A). Addition of preQ_1_ to either RNA (with or without Mg^2^+) shifted additional population to the high-FRET state while only slightly changing the *E*_*FRET*_ of either state.

**Figure 2.**
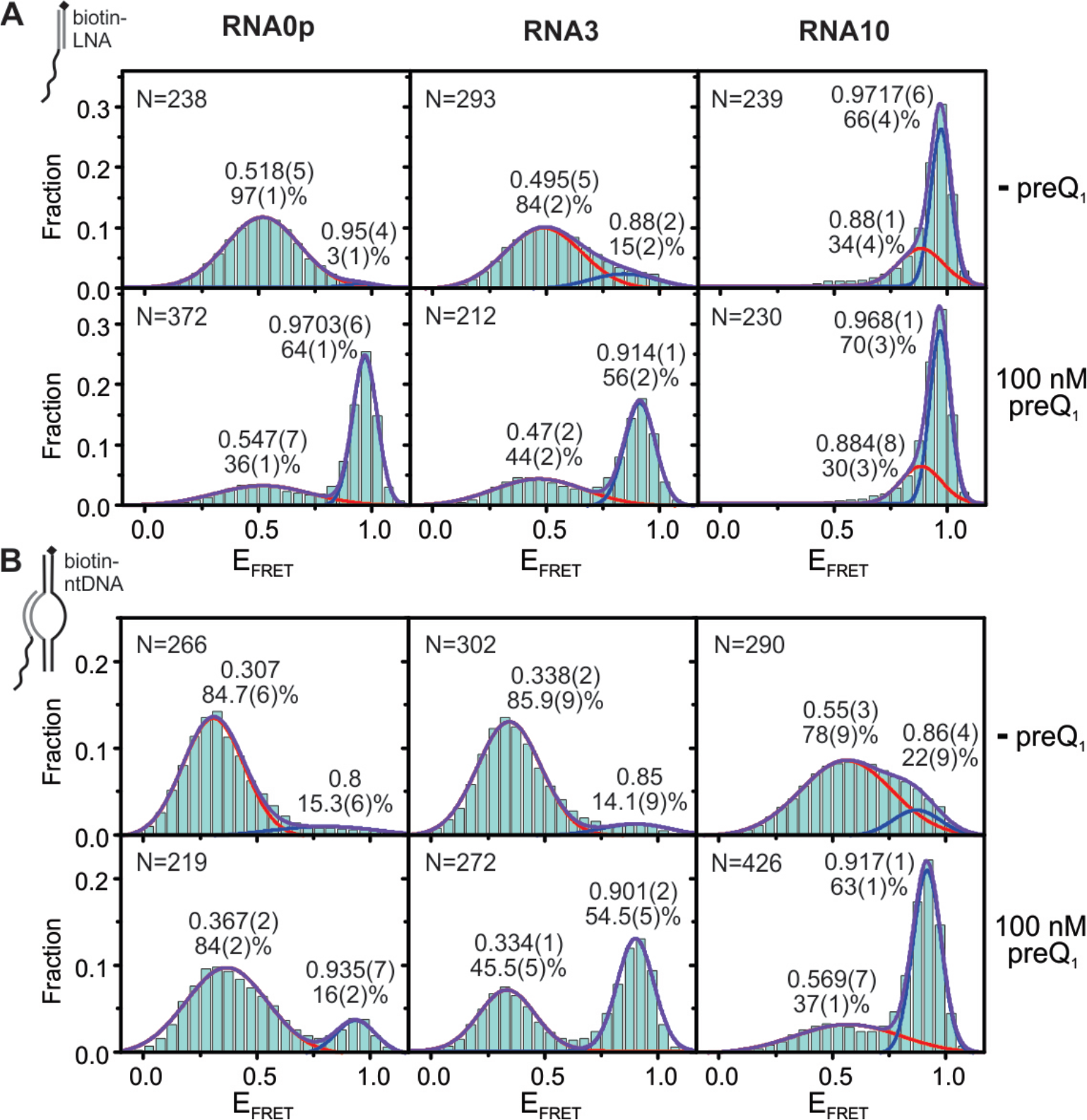
Riboswitch Folding Under Aptamer and Bubble Immobilization in the Absence of Mg^2+^; see also Figure S2. Each histogram was fit with two Gaussians, and the resulting *E*_*FRET*_ and population of each state are noted adjacent to the corresponding peak. The standard deviation of the last digit of each parameter (obtained via bootstrapping) is reported in parentheses. Values for which no standard deviation is reported were constrained during fitting. The number of traces included in each histogram (“N”) is noted in the upper left corner. (A) smFRET histograms of RNA0p (left), RNA3 (middle) and RNA10 (right) under aptamer immobilization in the absence of MgCl_2_ and the absence (top row) or presence (bottom row) of 100 nM preQ_1_. (B) Corresponding smFRET histograms under bubble immobilization.

In contrast, most RNA10 molecules occupied a very high-FRET state (*E*_*FRET*_ ~ 0.97) regardless of Mg^2+^ or preQ_1_ concentration (Figures 2A and 3A), consistent with an alternate structure. Specifically, additional nucleotides in RNA10 are predicted to form Watson-Crick base pairs with nucleotides that make up P2 and L2 in the native conformation (Figure S2B). To test this notion, we used mutagenesis to generate a sequence whose most stable predicted fold was once again the native structure. This mutant responded to ligand, confirming that when wild-type (WT) RNA10 is heat-annealed with CP, the native aptamer fold is disfavored.

To isolate the effects of the DNA template from those of RNAP, we next probed the riboswitch as part of a bubble complex that mimics the nucleic acid scaffold of an EC. Strikingly, addition of the DNA bubble led to a shift of population from the high-FRET to the mid-FRET state, and a shift of the mid-FRET state to lower *E*_*FRET*_ (~ 0.35 for RNA0p and RNA3, Figure 2B). Furthermore, the Mg^2+^-dependent shift of the mid FRET state to higher *E*_*FRET*_ was also greatly diminished (Figure 3B). RNA0p is rendered nearly insensitive to preQ1 and Mg^2+^, but sensitivity is restored as additional bases are inserted to form RNA3 and RNA10. Importantly, the effects of the bubble are relieved by increasing the K+ concentration (Figure S2C-D), suggesting that electrostatic repulsion by DNA interferes with preQ_1_-induced pseudoknot docking and Mg^2+^-induced compaction of the pre-docked conformation. Bubble-immobilized RNA10 properly responds to Mg^2+^ and preQ_1_, in contrast to its misfolding under aptamer immobilization, likely because the competing structure requires base pairs immediately adjacent to the bubble. These results show that the DNA template has the potential to play a significant, underappreciated role in shaping the folding free-energy landscape of nascent RNAs, as well as RNAs in the ubiquitous R-loops formed in other biological contexts (Richard and Manley, 2017).

**Figure 3.**
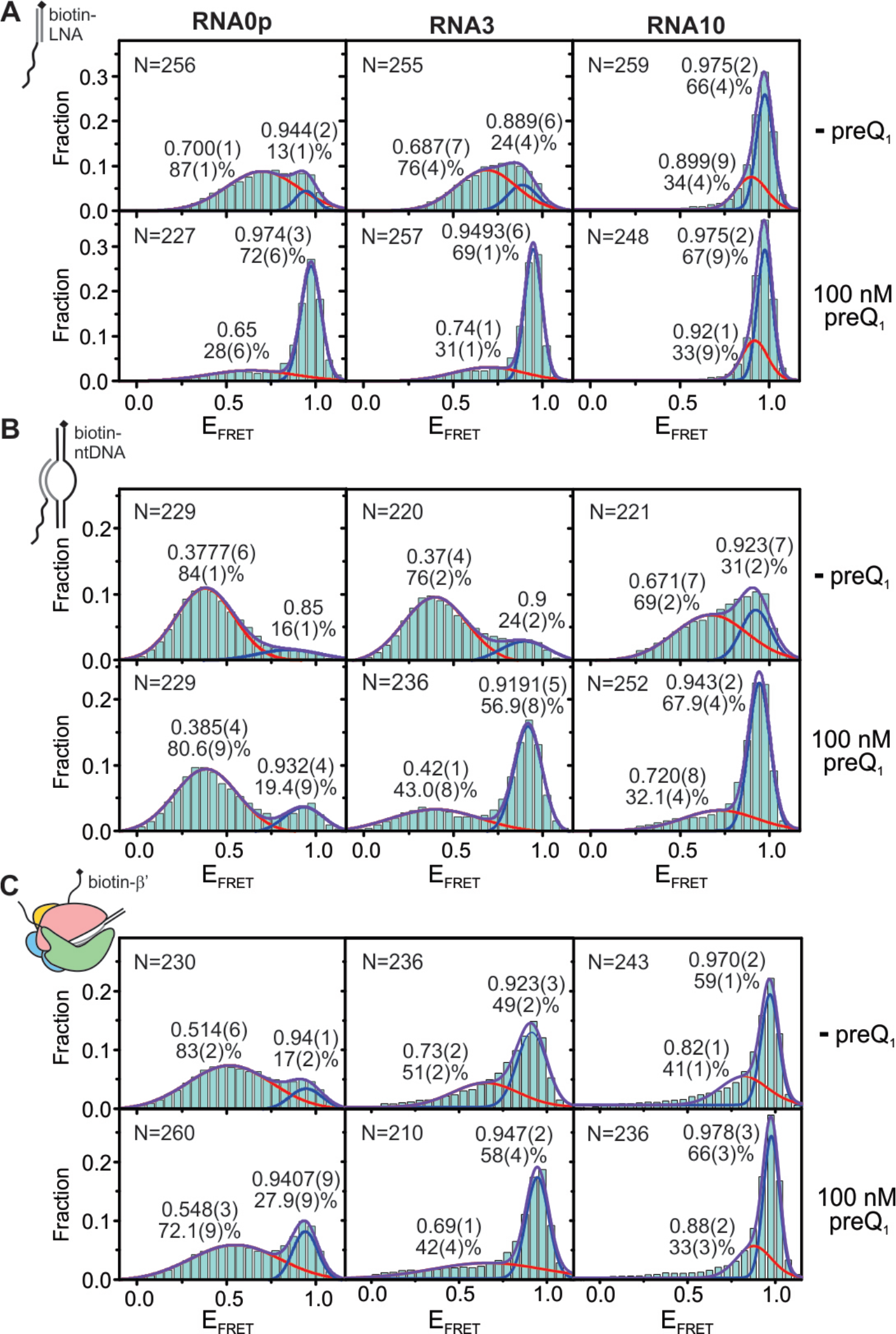
Riboswitch Folding Under Aptamer, Bubble and EC Immobilization in the Presence of 1 mM Mg^2+^; see also Figures S2 and S3. Each histogram was fit with two Gaussians, and the resulting *E*_*FRET*_ and population of each state are noted adjacent to the corresponding peak. The standard deviation of the last digit of each parameter (obtained via bootstrapping) is reported in parentheses. Values for which no standard deviation is reported were constrained during fitting. The number of traces included in each histogram (“N”) is noted in the upper left corner. (A) smFRET histograms of RNA0p (left), RNA3 (middle) and RNA10 (right) under aptamer immobilization in the presence of 1 mM MgCl_2_ and the absence (top row) or presence (bottom row) of 100 nM preQ_1_. (B) Corresponding smFRET histograms under bubble immobilization. Corresponding smFRET histograms under EC immobilization.

### Addition of RNAP to the Bubble Scaffold Restores Riboswitch Folding

To probe the direct effects of RNAP on riboswitch folding, we assembled active ECs on a bubble scaffold identical to that described above. We found the addition of RNAP to have an effect similar to that of increased salt, restoring favorable pseudoknot docking and compaction of the pre-docked conformation (Figures 3C and 4). Immobilization of ECs through biotinylated RNAP proved to be critical to proper quantification of these effects, as experiments with the biotinylated ntDNA revealed that ~50% of bubble scaffolds were not bound by RNAP (Figure 4A). Specifically, the FRET histogram of RNA3-containing ECs immobilized through biotinylated ntDNA can be fit as a linear combination of the bubble histogram and the RNAP-immobilized EC histogram, with the EC histogram contributing 51%. Consistent with this result, addition of GTP to ECs under identical conditions resulted in an elongation efficiency of 46% (Figure 4B). On-slide addition of the substrate nucleotide (GTP) to RNA0p-containing ECs caused the histogram of RNA0p to evolve toward that of RNA3, indicating that the ECs remain active on the slide (Fig. 4C). Notably, obtaining ECs active under imaging conditions required optimization of the oxygen scavenging system (OSS) used to minimize photobleaching and blinking (Aitken *et al.*, 2008) (Figure S3) to allow for stable binding of RNAP to the bubble scaffold and avoid nuclease degradation of the RNA.

**Figure 4.**
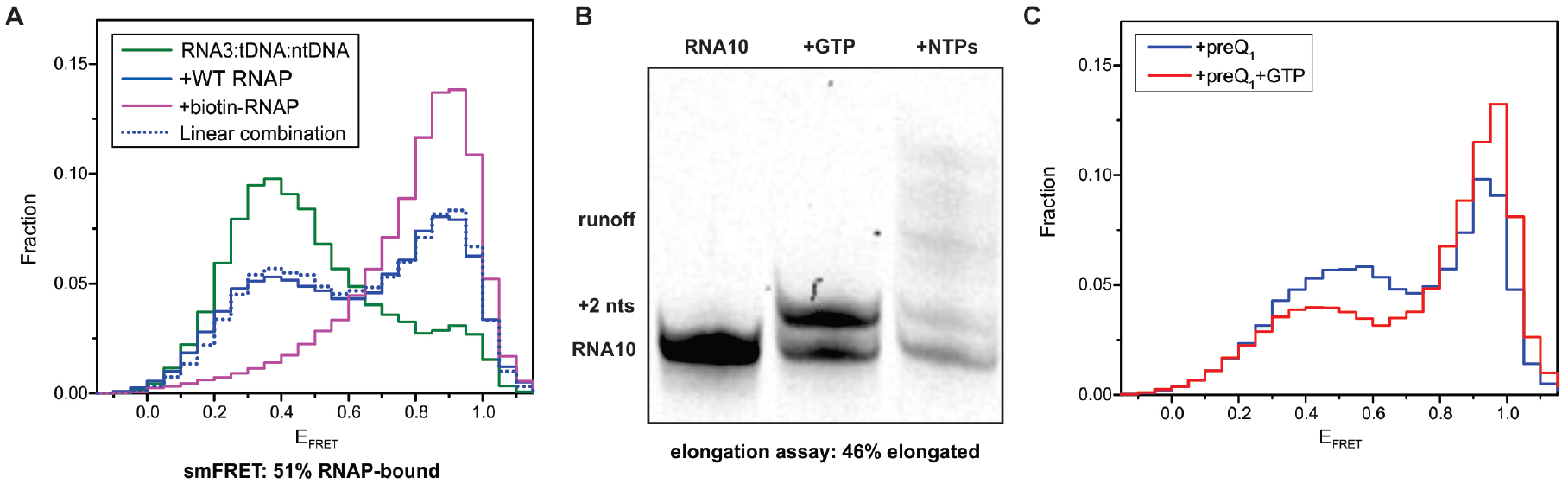
Assembly and Activity of Riboswitch-Containing ECs. (A) When RNA3-containing ECs are immobilized on a slide through biotinylated ntDNA, the FRET histogram (blue) corresponds to a linear combination (dashed blue) of the RNA3:tDNA:biotin-ntDNA histogram (green) and the RNA3:tDNA:ntDNA+biotin-RNAP histogram (purple), each contributing about 50%. (B) A representative gel from a bubble-initiated elongation assay (the gel that was quantified is shown in Figure S3A). (C) Incubation with GTP on the microscope slide causes the histogram of RNA0p ECs to evolve toward that of RNA3, indicating that nucleotides are being added.

Compared to bubble-immobilized complexes at the same low K^+^ concentration, the addition of RNAP shifted the mid-FRET state back to a higher *E*_*FRET*_ value (~0.53 for RNA0p), and shifted a significant fraction of population back to the high-FRET state (Figure 3C). Similar to bubble complexes, the closer the aptamer to the EC, the less stabilized the high-FRET state, which is evidence of steric hindrance. Surprisingly, however, the shift toward higher *E*_*FRET*_ occurs even for ECs containing RNA0p, in which the 3’ segment of the P2 helix is expected to be within the enzyme’s exit channel (Kang *et al.*, 2017).

To better rationalize these observations, we explored the steric accessibility of P2 folding using MD simulations with an all-heavy-atom structure-based model built on the cryo-EM structure of an *Eco* RNAP EC (Kang *et al.*, 2017), together with crystal (Klein *et al.*, 2009) and NMR (Kang *et al.*, 2009) structures of the preQ_1_ riboswitch (Figure 5). We found that while RNA0p is predicted to form fewer P2 base pairs than RNAs extended by 1-3 nucleotides (RNA1, RNA2 and RNA3), an average of 1-2 base pairs are intact (Table S3). In order for these base pairs to form, the 5’ segment of P2 must “bend back” toward RNAP. This bent-back pose places the 5’ leg of P2 near a cluster of basic residues in the β’ subunit and the flap-tip (FT) of the β subunit (Figures 5 and S4), conserved regions in the vicinity of the RNA exit channel (Guo *et al.*, 2018; Kang *et al.*, 2018).

**Figure 5.**
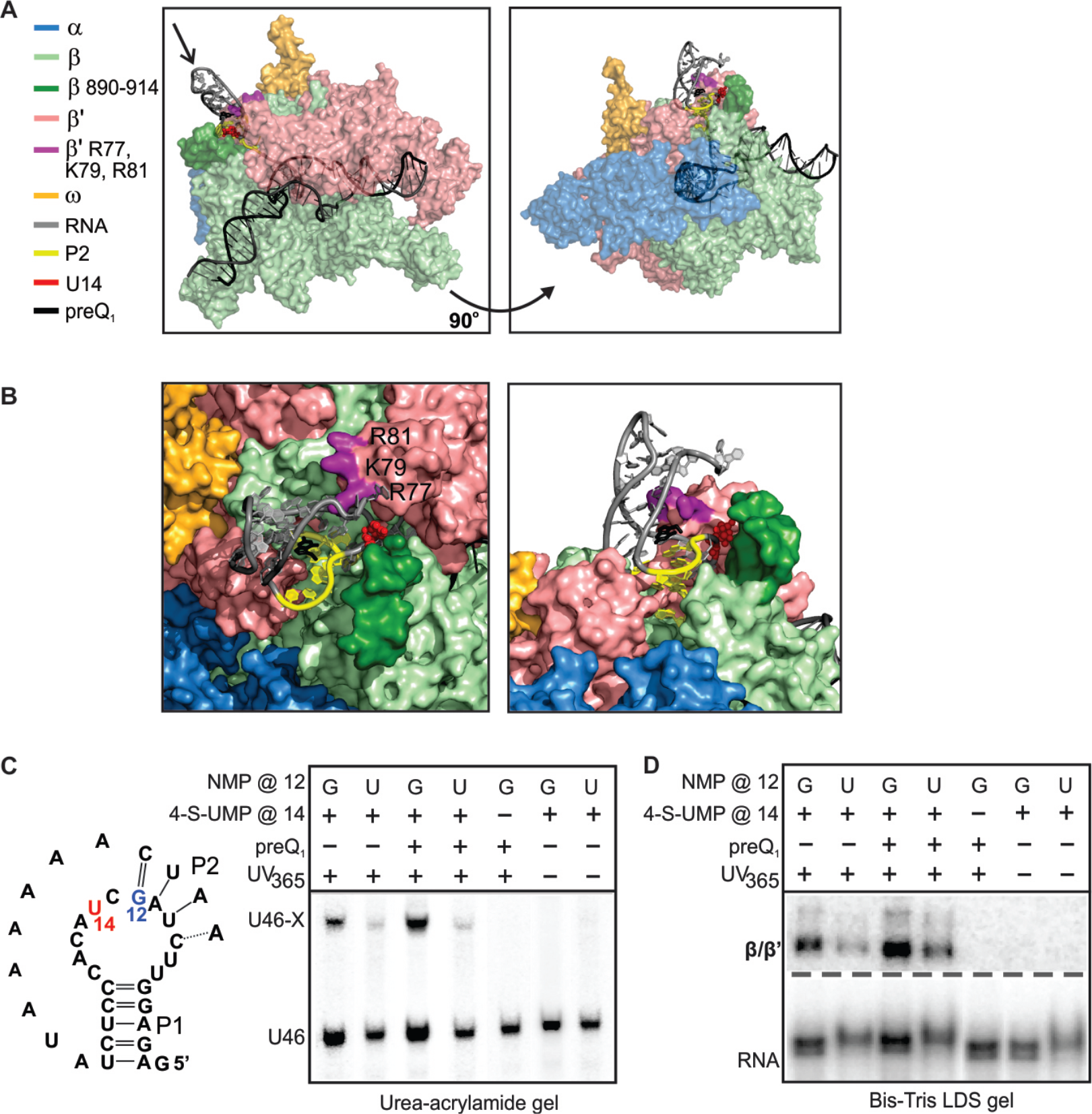
Simulations and Cross-Linking Experiments Show that P2 can Fold in the RNAP Exit Channel; see also Figure S4 and Table S3. (A) An MD simulation snapshot from two angles with RNAP subunits colored as in Figure 1 and P2 colored in yellow. The β flap-tip (residues 890-914) and β’ basic residues investigated later are shown in darker colors than the rest of their corresponding subunits. The model utilized a cryo-EM structure of an *Eco* EC (PDB ID: 6ALH), and crystal (PDB ID: 3FU2) and NMR (PDB ID: 2LIV) structures of the preQ_1_ riboswitch. (B) Left: Close-up views of the simulation snapshot in (A) as seen in the direction along the arrow (left), and from the same angle as the right-hand snapshot (right). (C) Left: schematic indicating position 12 within the aptamer, where a G-to-U mutation was investigated, and position 14, where a 4-thiouracil residue was incorporated for cross-linking. Right: Denaturing urea-acrylamide gel showing a shift of the RNA at the *que* pause upon cross-linking to RNAP, and controls lacking 4-thioU or UV illumination. (D) Bis-Tris LDS gel containing the same samples as panel (C), indicating that the RNA cross-links to β or β’. Experiments with an RNAP deletion mutant confirmed that the observed cross-links are to β’ (Figure S4E).

We probed for these predicted interactions through cross-linking experiments with 4-thio-UMP incorporated in or near P2. Indeed, when RNAP is stalled at the *que* pause site, both U10 and U14 efficiently cross-link to β’, confirming that, as predicted, the 5’ segment of P2 bends back to interact with RNAP (Figures 5 and S4). We found cross-linking to be dependent on the stability of P2, being more efficient in the presence of preQ1 and less efficient in a G12-to-U mutant, in which P2 is weakened by disrupting its only G-C base pair. Interactions between the A-rich tail L3 and the minor groove of P1 may also contribute to stabilization of the cross-linking competent conformation (Feng *et al.*, 2011; Klein *et al.*, 2009). In a complex stalled three nucleotides prior to the pause (C43, Figure S5B), we found cross-linking to be less efficient, presumably due to the inaccessibility of the 3’ segments of P2 and L3. Together, our simulations and crosslinking results indicate that, at the *que* pause, residues near P2 interact with the β’ subunit, potentially contributing to the stabilization of the docked conformation we observed through smFRET. Overall, our smFRET studies reveal that DNA and RNAP exert significant, opposing effects on nascent RNA folding, which are likely to play an important role in the regulation of transcription by other riboswitches.

### PreQ_1_ Releases a Pseudoknot-Stabilized Transcriptional Pause

Our results so far are consistent with a model in which the balance between electrostatic repulsion by the DNA template and stabilizing interactions with RNAP plays a significant role in preQ_1_- and Mg^2+^-dependent nascent RNA folding. Because RNAP pausing plays a critical role in guiding the co-transcriptional folding of numerous RNAs (Pan *et al.*, 1999; Perdrizet *et al.*, 2012), we next used single-round transcription assays to further probe the relationship between riboswitch folding and pausing. In addition to the *que* pause at U46, we observed a terminator that the riboswitch is expected to regulate (U70) and a second U-tract terminator further downstream (U108) (Figures 6 and S1). Nearly 100% of complexes paused at U46, with a pause half-life of 42 seconds in the absence of preQ_1_ and a vast majority population (97%) with a significantly shortened half-life of 15 seconds in the presence of preQ_1_. In addition, we found that the raw data set for *Bsu* provided by Larson et al. (Larson *et al.*, 2014) indicates that a transcriptional pause does, in fact, occur at this location *in vivo* (Figure S1E). Saturating concentrations of preQ_1_ increased termination efficiency modestly from 27% to 33%. Both effects can be fit globally with an apparent preQ_1_ binding affinity *K*_*1/2*_ of ~400 nM (Figure S5A), much weaker than the 5-10 nM for pseudoknot docking in the isolated aptamer, a reduction in apparent affinity also observed in other transcriptional riboswitches (Wickiser *et al.*, 2005). The effect of preQ_1_ on termination remains modest upon the addition of transcription factors, cell extract, crowding agents, additional riboswitch RNA, and varied NTP concentrations (Figure S5B-D), suggesting that *in vivo* non-canonical factors may be involved in regulation of the downstream genes (Baird *et al.*, 2010; Rinaldi *et al.*, 2016).

The effect of preQ_1_ and the similarity of the sequence around U46 to the pause consensus suggested that the *que* pause has both sequence and structural determinants, so to further uncover its origins, several RNA mutations were analyzed (Figures 6C-E and S6). Altering the C18 base that forms a Watson-Crick pair with preQ_1_ (Roth *et al.*, 2007) abolished the impact of preQ_1_ on pausing and termination, confirming that the riboswitch is responsible for the observed effects. A mutation that strengthens P2, stabilizing the docked state, (C9U, which converts a noncanonical C-A base pair to U-A) doubled the pause half-life (to 81 s), while the P2-weakening G12U mutation, which we also utilized in the cross-linking experiments presented above, shortened the pause (to 26 s) and eliminated the effect of preQ_1_. These aptamer mutations have smaller effects in the presence of preQ_1_, suggesting that the details of sequence-and structure-dependent interactions with RNAP are more critical in the less-folded *apo* state. As expected, mutation of key bases in the pause consensus sequence (G37U and G36U/G37U) greatly reduced pause half-life in the absence of preQ_1_ to 12 and 10 s, respectively. Addition of one U residue between the aptamer and the pause site (1U) also weakened pausing in the absence of preQ_1_ (23 s), with little effect in the presence of preQ_1_, and adding a second U (2U) had no significant further effect (22 s). The position of the pause is dictated by the consensus sequence rather than the riboswitch, as these insertions shift the pause site downstream by one and two nucleotides, respectively. These results indicate that the consensus sequence and strongly distance- and conformation-dependent interactions between the riboswitch and RNAP cooperate to stabilize the paused state, with ligand binding stabilizing the pseudoknot in a distinct way that facilitates pause release.

### Interactions with the β Flap-Tip of RNAP Stabilize Pausing

We next tested the effects of RNAP-riboswitch contacts revealed by MD simulations. We first substituted basic β’ residues (R77A, K79A and R81A, Figure 5B) that lie near the stem of the *his* pause hairpin in *E. coli* ECs (Guo *et al.*, 2018; Kang *et al.*, 2018) and have been implicated in the function of *put* antiterminator RNA structures (Sen *et al.*, 2002). Arg77 is conserved between *Eco* and *Bsu* RNAPs, and Lys79 and Arg81 are also positively charged in *Bsu* RNAP. We found that individual point mutations led to modest decreases in pause half-life in the absence of preQ_1_, whereas a triple mutant had the most significant effect (~2-fold). In contrast, effects on pausing were negligible in the presence of preQ_1_ (Figure S6B). This suggests that the pause is partially stabilized by electrostatic interactions between the exit channel and the pause-promoting, ligand-free docked conformation of the riboswitch. All variants showed decreased termination at U70, with the R77A and triple mutants showing the most extreme effects, though the termination-enhancing activity of preQ_1_ was retained (Figure S5E and I).

We next tested the effect of the β FT, which is required for the pause-stabilizing effects of RNA duplexes (Hein *et al.*, 2014). In contrast to the point mutants, which had effects only in the absence of preQ_1_, deleting the FT by removing residues K890-K914 of β (ΔFT) decreased the pause half-life by about 30% in both the presence and absence of preQ_1_ (Figure S6C). The effect of preQ_1_-dependent conformational changes on pausing was therefore maintained for ΔFT RNAP, despite an overall decrease in pause half-life. In contrast, we observed no effect of the G12U mutation on pausing by ΔFT RNAP, indicating that pausing by ΔFT is not stabilized by the pseudoknot. Together, our experiments on RNAP variants suggest that both electrostatic interactions with exit channel residues and steric interactions with the FT contribute to pseudoknot-dependent pause stabilization in the absence of preQ_1_. By contrast, in the preQ_1_-bound docked state, only steric interactions with the FT contribute to pause destabilization. These observations support the notion that the docked states in the presence and absence of preQ_1_ are distinct, and involve different interactions with RNAP.

## DISCUSSION

We used smFRET, biochemical transcription assays and MD simulations to study the co-transcriptional folding of the *Bsu* preQ_1_ riboswitch. Isolated aptamers exhibited the previously observed pre-docked (mid-FRET) and docked (high-FRET) conformations, with the docked conformation being stabilized upon addition of preQ1 and Mg^2+^, and the pre-docked conformation becoming more compact upon addition of Mg^2^+. Similar Mg^2+^-dependent changes in the absence of ligand have previously been observed by NMR in the isolated *Bsu* and *Fusobacterium nucleatum (Fnu)* preQ_1_ riboswitches. In the *Bsu* riboswitch, addition of Mg^2+^ leads to changes in the ligand-binding pocket that make it resemble but not perfectly mimic the ligand-bound state (Suddala *et al.*, 2013). In the *Fnu* riboswitch, addition of Mg^2+^ leads to signals suggestive of P2 base pairing and interactions between P1 and the 3’ tail (Santner *et al.*, 2012). These structural insights may explain some of the variation in *E*_*FRET*_ that we observe upon addition of Mg^2^+ and ligand. For example, we found that addition of preQ_1_ shifts the *E*_*FRET*_ of the docked conformation to slightly higher values, a change that accompanies the concomitant rearrangement of the ligand-binding pocket. The significant changes in the *E*_*FRET*_ of the pre-docked conformation upon addition of Mg^2+^ are consistent with formation of the P1-3’ tail interactions observed in both the *Bsu* (Suddala *et al.*, 2013) and *Fnu* (Santner *et al.*, 2012) preQ_1_ riboswitches.

We also uncovered a previously undescribed effect of the DNA template that disfavors nascent RNA folding in a distance-dependent manner. The *E*_*FRET*_ of the predocked conformation was significantly lowered and its dependence on Mg^2+^ weakened by addition of the DNA. We found that addition of KCl increased the *E*_*FRET*_ of the predocked conformation in a continuous fashion (Figure S3C-D), suggesting that there is no one distinct pre-docked conformation or that its dynamics remain unresolved. The addition of RNAP reverses this effect, restoring preQ1-dependent docking and compaction of the pre-docked state despite steric hindrance of folding in the exit channel (Figure 3). MD simulations and cross-linking experiments provide additional support for the notion that P2 can fold in the exit channel, showing that a subset of P2 base pairs are intact even in RNA0p, and that at the *que* pause, the 5’ segment of P2 bends back to interact with RNAP residues near the exit tunnel (Figure 5). Our observations provide concrete evidence for a previously-hypothesized chaperone activity of RNAP, which has proven difficult to investigate by other means (Pan and Sosnick, 2006; Zhang and Landick, 2016).

Few studies exist in which naked RNA was directly compared to RNA in bubble complexes or ECs, with the only changes being the addition of those components of the transcription machinery. A study utilizing the fluorescent adenine analogue 2-aminopurine showed that addition of the DNA template had little effect on base stacking in an RNA hairpin, while addition of bacteriophage T7 RNAP to the bubble decreased stacking significantly (Datta and Hippel, 2008). A similar pattern was observed with single-stranded nascent RNA (Datta *et al.*, 2006). Our investigation reveals very different behavior for a pseudoknot, with the DNA template having a significant effect and RNAP favoring RNA folding. Addition of *Eco* RNAP to a bubble scaffold decreased the rate of annealing of an antisense oligonucleotide (designed to mimic folding of the *his* pause hairpin) to the nascent RNA, but to a lesser extent than might be expected, indirectly implying a chaperoning function of RNAP (Hein *et al.*, 2014). However, Hein et al. investigated a bimolecular annealing process, and their results therefore cannot be directly compared to our measurements of intramolecular pseudoknot folding. Uhm et al. used smFRET to compare thermally annealed naked RNA to RNA that was transcribed on the slide surface by T7 RNAP. They found that the TPP riboswitch was more sensitive to ligand when it had been transcribed, and that the location at which RNAP was stalled greatly affected ligand sensitivity (Uhm *et al.*, 2018). While these results could be attributed to direct interactions between the riboswitch and RNAP, the kinetic effects of folding during transcription cannot be ruled out as the predominant factor.

Conversely, using transcription assays, we showed that riboswitch folding modulates RNAP pausing (Figure 6). Current evidence points to the existence of an “elemental” paused state, in which sequences similar to the consensus trigger conformational changes that interfere with nucleotide addition. Elemental pauses have so far been found to be stabilized by two primary mechanisms (Artsimovitch and Landick, 2000; Zhang and Landick, 2016) (Figure 7A). In “hairpin-stabilized” or Class I pausing, a hairpin in the nascent RNA causes conformational rearrangements in the exit channel that stabilize the pause by allosterically promoting a “swiveled” conformation of RNAP (Kang *et al.*, 2018). Class I pauses are further stabilized by the transcription factor NusA through interactions with the nascent RNA and the β flap domain (Hein *et al.*, 2014; Toulokhonov *et al.*, 2001), and are resolved by the trailing ribosome (Landick *et al.*, 1985). In a second mechanism (Class II), the elemental pause enables backtracking by RNAP, leading to disengagement of the 3’ end of the RNA from the active site. Class II pauses are sensitive to GreB, which enhances RNA cleavage by RNAP, and NusG, which antagonizes backtracking (Herbert *et al.*, 2010). The Class II *ops* pause provides the opportunity for the antitermination factor RfaH to bind, which temporarily stabilizes the pause before promoting elongation and suppressing pausing elsewhere (Artsimovitch and Landick, 2002).

**Figure 6.**
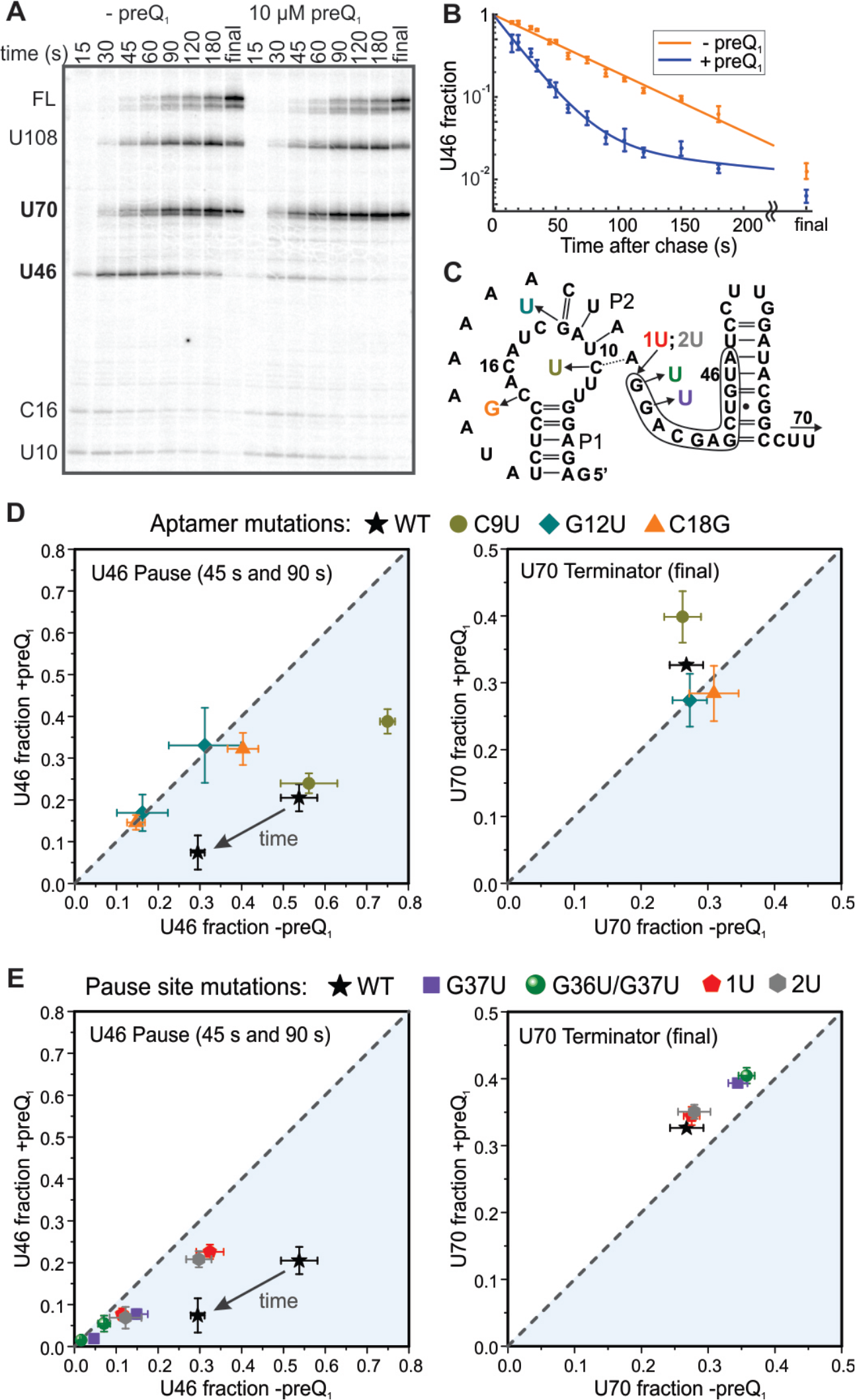
Effects of preQ_1_ on Transcriptional Pausing and Termination; see also Figures S5 and S6 and Tables S4 and S5. All error bars represent the standard deviation of three independent replicates. (A) Representative denaturing urea-acrylamide gel close-up showing two weak pauses within the aptamer domain (U12 and C16), the *que* pause immediately following the aptamer (U46), the terminator regulated by the riboswitch (U70), a U-tract arrest site (U108) and full-length RNA (FL). Here and elsewhere, all lanes and data points marked “final” were recorded after a 3-5 minute incubation with 200 μM NTPs. (B) Fraction of complexes at the U46 site as a function of reaction time. A single-exponential fit is shown for the time course in the absence of preQ_1_ (orange), and a double-exponential fit is shown for the time course in the presence of 10 μM preQ_1_ (blue). (C) Sequence of the aptamer and expression platform showing the final nucleotide added at the bands indicated in panel A, as well as the color-coded locations of mutations reported in panels D and E. U70 and U108 are beyond the region shown. The consensuslike pause sequence is outlined. (D) Effects on pausing at U46 (left) and termination at U70 (right) of mutations within the aptamer domain. Data points above the dashed line indicate that addition of preQ1 increased the population of a particular species, while points below the dashed line indicate that preQ_1_ decreased the population of that species. PreQ_1_ had no effect on data points that fall on the dashed line. (E) Effects on pausing (left) and termination (right) of mutations at the pause site.

The *que* pause we studied similarly requires an initial elemental pause, evidenced by the decrease in pause lifetime upon mutation of the consensus sequence (G37U and G36U/G37U mutants; Figures 6E and S7). The *que* pause is destabilized by moving the riboswitch further from RNAP (1U and 2U insertions), destabilizing P2 (G12U mutant), or deleting the β FT. We propose that these variants represent elemental or only weakly pseudoknot-stabilized pauses, lasting 22-25 seconds under our experimental conditions (Figure 7). In the WT and C9U variants, in contrast, the elemental pause is further stabilized by interactions between the pseudoknot and RNAP, resulting in a 2- to 4-fold increase in the pause half-life over that of the elemental pause.

**Figure 7.**
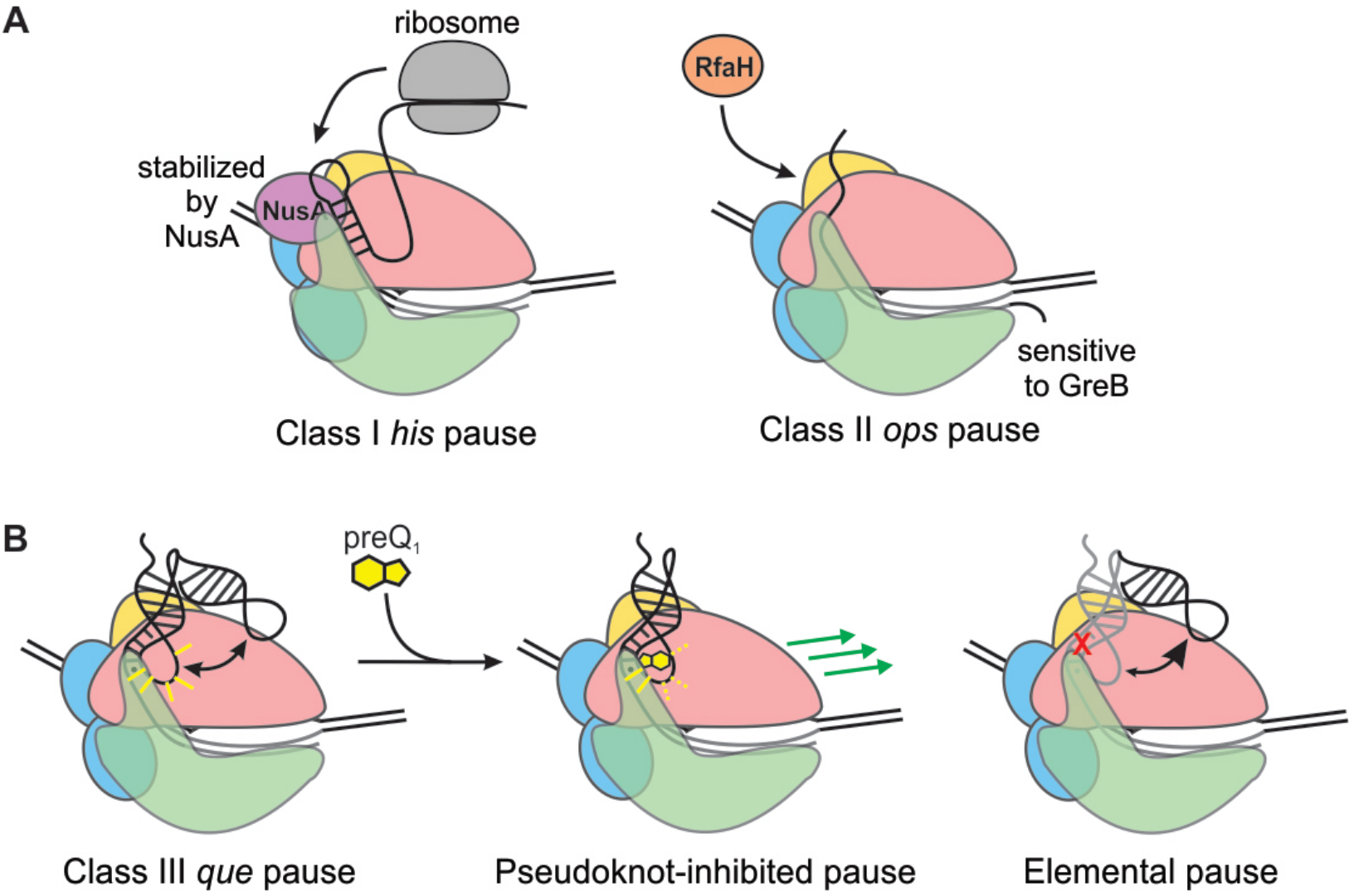
Mechanisms for Pause Stabilization and Control of Gene Expression; see also Figure S7. (A) At a Class I pause, the trailing ribosome promotes RNAP escape and couples transcription to translation to regulate attenuation in amino acid biosynthesis operons. At a Class II pause, RfaH binds to the EC and antiterminates transcription of long operons. (B) At the Class III *que* pause, the preQ_1_ ligand chases RNAP from the pause and rearranges the riboswitch structure to promote termination, converting a pseudoknot-stabilized pause (left) into a pseudoknot-inhibited pause (middle). Disrupting the pseudoknot yields an elemental pause (right).

These results suggest an analogy to the hairpin-stabilized Class I pause, and the *que* pause described here indeed shares intriguing characteristics with other pauses that are dependent on nascent RNA structure. Notably, the first base-pair of P2 is the same distance from the 3’ end of the nascent RNA as is the first base-pair of the *his* pause hairpin (Toulokhonov *et al.*, 2001). This is also identical to the distance at which initial folding of a pseudoknot was observed in the fluoride riboswitch (Watters *et al.*, 2016). Differences in RNAP translocation register yield a 1-2 nucleotide variation in the position of the paired base relative to RNAP; nevertheless, these similarities indicate a common length-scale for nucleation of RNA folding. Insertion of two nucleotides between the hairpin and the *his* pause site reduces the pause lifetime and eliminates the ability of an oligonucleotide that disrupts the hairpin to inhibit pausing (Toulokhonov *et al.*, 2001), suggesting that the hairpin no longer affects pausing at this distance. Our 1U and 2U variants exhibit the same effect, with potential modulation of nascent RNA structure through preQ_1_ binding having little effect on pausing. Similar to our results, it has previously been found that deletion of the β FT has no effect on the duration of an elemental pause, while it destabilizes a *his* pause-complex mimic in which the hairpin is replaced with a duplex (Hein *et al.*, 2014). The *que* pause therefore shares certain characteristics with the Class I *his* pause despite being stabilized by a pseudoknot rather than a hairpin.

However, there are several significant differences between the properties of the *que* pause and canonical Class I and Class II pauses. The FT has a much larger effect on Class I pauses than on the *que* pause, and the presence of a pseudoknot rather than a hairpin also renders the *que* pause insensitive to NusA (Figure S5G and S6D), which requires either a simple duplex of at least 10 base-pairs (Kolb *et al.*, 2014) or a loop of greater than 4 nucleotides (Toulokhonov *et al.*, 2001) for maximum stimulation of pausing. The folded riboswitch instead presents a 4 base-pair helix at the same position as the *his* pause hairpin, and instead of a loop presents a triplex-like structure in which L3 forms Hoogsteen-face interactions with the helix P1 (Klein *et al.*, 2009). The riboswitch therefore lacks the structural features that confer NusA sensitivity in hairpin-dependent pauses. PreQi binding has the effect of chasing RNAP away from the *que* pause, a function performed by the ribosome in the case of Class I pauses (Landick *et al.*, 1985). Unlike Class II pauses, the RNA at the *que* pause does not undergo cleavage in response to GreB (Figure S5H), indicating that RNAP is not in a backtracked position. Our results suggest that the *que* pause represents a new mechanism in which an elemental pause is stabilized by a pseudoknot in the nascent RNA next to the RNAP exit channel, which we term “pseudoknot-stabilized” or “Class III” pausing (Figure 7B).

Our cross-linking results would seem to indicate that addition of preQ_1_ strengthens interactions with RNAP (Figure 5), and on the basis of our transcription assays in the absence of preQ_1_, this would be expected to promote pausing. However, in our cross-linking experiments, pause release is impossible due to the absence of NTPs. In this situation, preQ_1_ has the sole effect of stabilizing a bent-back, cross-linking competent conformation in which P2 is intact. When the presence of NTPs makes pause release possible, this conformation, which clearly differs from the pause-promoting bent-back conformation, appears to instead yield “pseudoknot-inhibited” analogue of the hairpin-inhibited pause escape mechanism (Zhang and Landick, 2016). The G12U mutant does not transition into a pseudoknot-inhibited pause because it binds preQ1 too weakly, leading to minimal effects of preQ_1_ on pausing and termination. The 1U and 2U variants do not transition into a pseudoknot-inhibited state for one or both of two reasons: 1) The increased distance between the pseudoknot and RNAP means that RNAP does not “feel” the conformational change upon preQ_1_ binding; and/or 2) preQ_1_ binding does not cause an extensive structural rearrangement, as seen in our MD and smFRET results.

In summary, we have demonstrated an intimate cross-coupling between bacterial riboswitch folding and transcriptional pausing that is likely to govern the co-transcriptional folding of numerous RNAs. Using a combination of approaches that are readily applicable to a wide variety of RNA species, we found that the presence of stalled RNAP provides a favorable free energy landscape for folding of the nascent preQ_1_ riboswitch. In turn, the ligand-free but pre-folded riboswitch pseudoknot stabilizes the paused state of RNAP via interactions with the β FT and with positively charged residues on the β’ subunit, while preQ_1_ binding stabilizes a distinct docked conformation that counteracts pausing. As pseudoknots are a common RNA structural motif, we expect that numerous examples of Class III pausing will be found among the thousands of transcriptional pauses that have been identified (Larson *et al.*, 2014; Vvedenskaya *et al.*, 2014). Our results reveal an additional, so far underappreciated layer to bacterial gene regulation that can be exploited for antibiotic drug intervention (Blount and Breaker, 2006; Deigan and Ferré-D’Amaré, 2011; Howe *et al.*, 2015).

## SUPPLEMENTAL INFORMATION

Supplemental information includes seven figures and six tables.

## AUTHOR CONTRIBUTIONS

Conceptualization, J.R.W. and N.G.W.; Methodology, J.R.W., Y.A.N., R.L.H.; Investigation, J.R.W., Y.A.N., V.R., R.L.H.; Writing - Original Draft, J.R.W.; Writing - Review & Editing, J.R.W., Y.A.N., V.R., R.L.H., C.L.B. III, I.A., N.G.W.; Supervision, N.G.W., I.A., C.L.B. III; Funding Acquisition, N.G.W., I.A., J.R.W.

## ACKNOWLEDGEMENTS

We thank Dr. Catherine Eichhorn and Dr. Hashim Al-Hashimi for generous gifts of plasmids containing the preQ_1_ riboswitch, Dr. Paul Lund for help mining previously published deep sequencing data, and Dr. Tina Henkin for a gift of *Bsu* RNAP. This work was supported by National Institutes of Health (NIH) R01 grants GM062357 and GM118524 to N.G.W., R01 grant GM67153 to I.A., R01 grant GM037554 to C.L.B. III, and NIH NSRA F32 postdoctoral fellowship GM113297 and K99 pathway to independence award GM120457 to J.R.W. V.R. acknowledges the Undergraduate Research Opportunity Program and Department of Biophysics for additional funding.

